# Deep learning pipeline reveals key moments in human embryonic development predictive of live birth in IVF

**DOI:** 10.1101/2023.06.27.546684

**Authors:** Camilla Mapstone, Helen Hunter, Daniel Brison, Julia Handl, Berenika Plusa

**Affiliations:** Faculty of Biology, Medicine and Health (FBMH), Division of Developmental Biology & Medicine, Michael Smith Building, University of Manchester, Manchester, United Kingdom; Alliance Manchester Business School, University of Manchester, Manchester M15 6PB, UK; Department of Reproductive Medicine, Old Saint Mary’s Hospital, Manchester University NHS Foundation Trust, Manchester Academic Health Science Centre, Manchester, UK; Division of Developmental Biology and Medicine, Maternal and Fetal Health Research Centre, School of Medical Sciences, Faculty of Biology Medicine and Health, University of Manchester, Manchester Academic Health Science Centre, Manchester, UK

## Abstract

Demand for IVF treatment is growing, however success rates remain low partly due to difficulty in selecting the best embryo to be transferred. Current manual assessments are subjective and may not take advantage of the most informative moments in embryo development. Here, we apply convolutional neural networks (CNNs) to identify key windows in preimplantation human development that can be linked to embryo viability and are therefore suitable for the early grading of IVF embryos. We show how machine learning models trained at these developmental time-points can be used to refine overall embryo viability assessment. Exploiting the well-known capabilities of transfer learning, we illustrate the performance of CNN models for very limited data sets, paving the way for the use on a clinic-by-clinic basis, catering for local data heterogeneity.

## Main Text

Machine learning (ML) has already demonstrated great promise in many areas of medical imaging^1-3^ and has the potential to revolutionise the field of medical diagnostics. There has been recent interest in applying ML approaches to tackle infertility, a growing health crisis impacting both individuals and society that has led to a rising demand for in vitro fertilisation (IVF) treatments^4^. Due to low success rates multiple IVF attempts are often required, leading to additional cost and distress for the patients. One of the main challenges in IVF is the difficulty in selecting an embryo to be transferred, as the general consensus on what the healthy, human embryo looks like, and which embryos are the most suitable for transfer is still under extensive debate.^5-7^ The selection process routine in most clinics involves visual assessment of embryos in real time or via time-lapse videos. Embryos are assessed based on morphological features such as the number and size of cells at cleavage stage and in the trophectoderm (TE) and inner cell mass (ICM) in the blastocyst (Fig. 1A), the expansion of the cavity, developmental timings, cellular fragmentation, and multi-nucleation. This manual assessment is subjective, with up to 83% variation between embryologists^8^, and uses only a fraction of the information potentially available. A convolutional neural network (CNN) based approach that automatically assesses embryos using more extensive information from across the time-lapse videos could potentially provide a consistent and reproducible method for embryo selection^9^.

**Figure 1:**
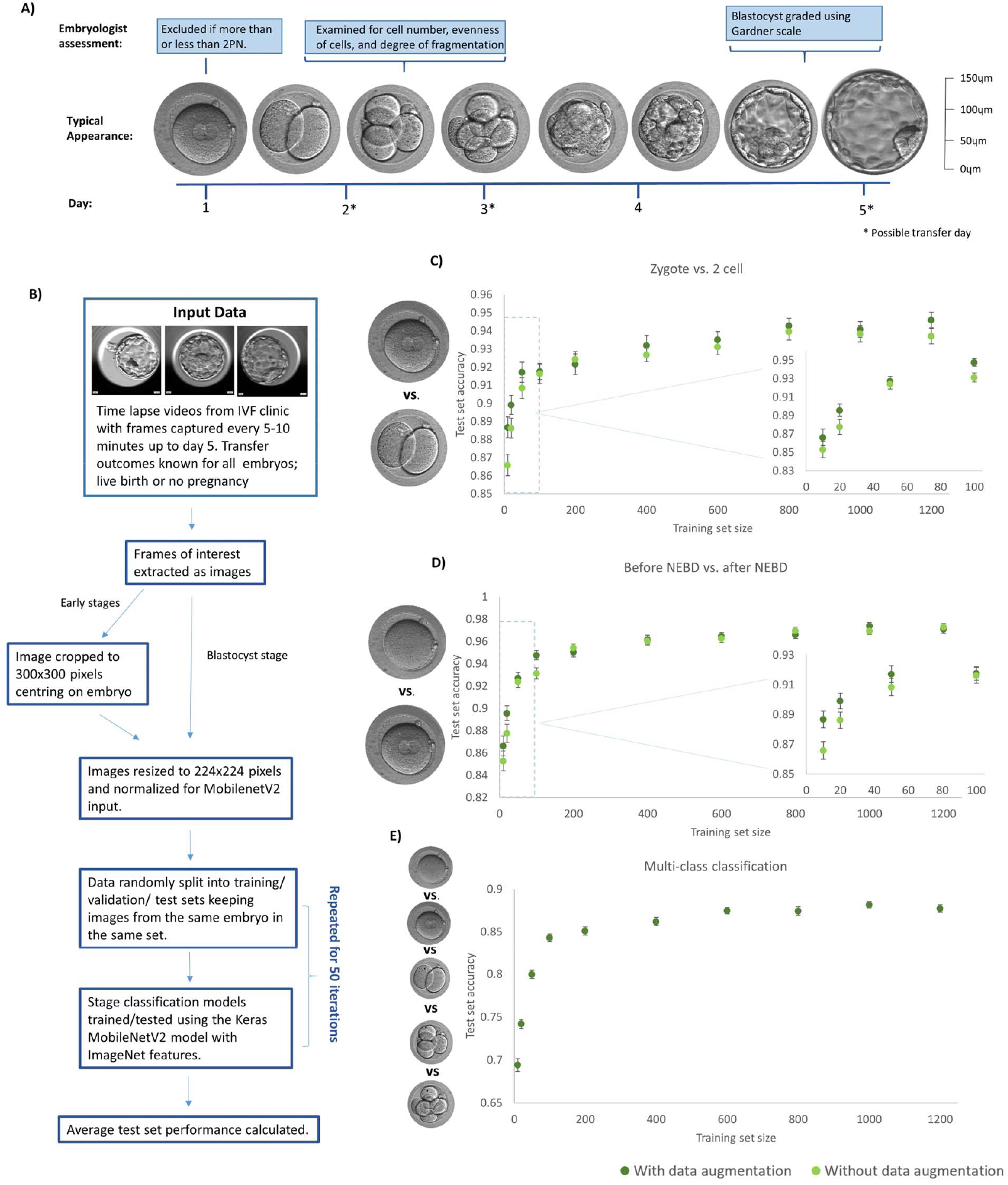
Performance of developmental stage classification models with increasing amount of training data. A) The typical development of human embryo and an overview of embryologist assessments carried out at each stage. B) The methodology of training developmental stage classification models from time-lapse data. C) The accuracy of the test set on models trained to classify an image as zygote or 2 cell with varying amounts of training data. D) The accuracy of the test set on models trained to classify an image as before NEBD or after NEBD with varying amounts of training data. E) The accuracy of the test set on multi-class models trained to classify an image into one out of 5 developmental stage classes with varying amounts of training data. C-E) all show the average scores over 50 training attempts with 25 embryos in each class for the test and validation set. The accompanying error bars are the standard error across these 50 training attempts.

In recent years many CNN approaches to assessing embryo viability have been developed^8,10-23^, demonstrating the potential for deep learning to inform embryo selection. Most of these CNN-based studies focus on the blastocyst stage, mirroring the most widely used process of embryo selection by the trained embryologists. Although embryologists may pay attention to earlier stages, there is no consensus on which stages are crucial to look at and no standardised method of combining morphological features and/or assigned grades from across development into a single overall viability score. A few CNN based approaches have incorporated frames from earlier embryonic stages^12,15,17,22,23^, and some have succeeded in making viability assessments using solely pre-blastocyst stages^15,17,23^, demonstrating that it is possible for CNNs to extract information about embryo viability from this early developmental period. It is yet to be determined whether there are specific moments in this period that are particularly informative, therefore the choice of which frames to include requires further investigation. This choice could impact model performance, as frame selection based just on video timestamp could miss crucial moments or add unnecessary data, potentially leading to overfitting. Identifying moments in development most predictive of live birth could lead towards further improvements in viability assessment, based on the most important information across development.

In this work, we used a CNN to determine the moments in development that can be linked to higher embryo viability (defined as the most predictive of life birth) and demonstrated the importance of including these developmental time-points when assessing embryo viability. Firstly, through development of a stage classification algorithm we demonstrated the high efficiency of our chosen model at correctly classifying stages of pre-implantation embryos. We then showed that this ML framework can assess blastocyst viability at a similar level to highly trained embryologists. Subsequently, we successfully applied this model to predict live birth at earlier stages, for which no standardised assessment methods exist. Capitalising on that, we used ML to identify previously unknown pivotal time-points for live birth prediction in the pre-blastocyst embryo, which is a first step for better understanding of the biology of human development and offers the means for better overall viability assessment and/or earlier transfer. Finally, we showed that combining information from our identified time-points of embryo development with blastocyst stage predictions allows for a quantitative ranking of high-quality blastocysts which could not have been achieved by any of the existing assessment methods. Critical to widespread clinical application, our findings also confirm that it is possible to train a CNN model in this domain to a high standard using limited data.

## Results

In this study we employed CNNs to analyse the biological process of human pre-implantation development and provide information of clinical relevance. To this end, we trained and tested CNN models on a dataset of time-lapse videos of embryos (700 in total) with known transfer outcomes from the IVF clinic at the Department of Reproductive Medicine, St Marys’ hospital, Manchester, UK. This is an NHS-funded clinic with stable patient population demographics and standardised treatment policies, and therefore highly suitable for single centre treatment outcome studies^24^. Embryos were cultured in either Embryoscope ™ or Embryoscope+ ™ time-lapse incubator system (Vitrolife, Sweden).

Live birth prediction is a difficult task as some embryos fail due to maternal factors rather than defects of the embryo itself. This essentially adds ‘label noise’ to the data; some ‘unsuccessful’ embryos will actually be of a high quality and likely developmentally competent. Therefore, in addition to predicting live birth, we first trained and tested models on the simpler task of developmental stage classification in order to fully evaluate the capabilities of our ML framework on this dataset.

### Developmental stage classification

To develop stage classification models, all time-lapse videos had one frame manually extracted for each developmental stage, resulting in equal class sizes. To reduce the amount of training data needed, we used the established practice of transfer learning, using the MobileNetV2 model with layers pre-trained on ImageNet, the overall workflow is illustrated in Fig. 1B.

First, we trained a binary model to classify embryos as zygote or 2-cell stage (Fig.1C). Subsequently, we trained a second binary model to recognise subtler differences on the subcellular level. To this end, we used images of embryos taken before or after NEBD (Fig.1D). The two investigated models were trained with varying amounts of training data by randomly excluding a portion of our data set. The training set size ranged from 10 to 1200 (all data used) with and without data augmentation. The validation and test sets were kept constant at 100 images each. The average test set scores (Fig. 1C-D) showed that when all the data was used, we reached a test set accuracy of 97.1% for the zygote vs 2-cell model and 94.6% for the before NEBD vs. after NEBD model.

For both models the accuracy increased with the amount of training data, as expected, and appeared to be reaching a plateau in performance once the number of images in the training set rose to around 200-400. This plateau may suggest that the model performed close to its optimal level and we are not likely to gain much better performance by adding more training data. Visual inspection of the images (Fig. s1A) also suggested that there is an upper limit to the accuracy that could be obtained as vacuoles and cells dividing in a plane that does not allow subsequent blastomeres to be immediately identified can cause an embryo to appear to be at a different stage when viewed without the context of the time-lapse video.

Our results also confirmed that relatively high performances can be achieved even with very little data. Over 85% test set accuracy was obtained for both models with a training set size of just 10 (5 images from each stage). Fig. 1C-D also show that augmentation (see methods for details) seemed to have a small but positive effect, especially with limited training data, so we chose to continue using augmentation going forward.

We then trained multi-class models to classify images into five output classes corresponding to the consecutive stages of human pre-implantation development (Fig. 1E). We saw that a test set accuracy of 87.7% was achieved, which is much higher than the by-chance score of 20%. The results in Fig. 1E show a similar trend to the one observed in the binary models; increases in accuracy became small once the training set reached about 200-400 images, and a reasonably high test set accuracy (69.4%) was achieved even with a training set of just 10. These results further confirm that MobileNetV2 with pre-trained layers can achieve a high performance, even with a small amount of the training data, when analysing routinely collected images of preimplantation human embryos.

### Live birth prediction at blastocyst stage

Following the successful development of developmental stage classifiers, we next used the same model framework (MobileNetV2 with transfer learning via layers pre-trained on ImageNet) to predict live birth based on the blastocyst stage, as this stage is most commonly used for embryo assessment so is a good time-point for comparisons. We extracted the final frame from each time-lapse video to produce a dataset of blastocyst images and trained the model on this dataset using five-fold cross validation with 50 repeat training runs for each fold. For each embryo in the dataset, we calculated an average model confidence score from all 50 training runs where that embryo was in the test set. We then calculated the ROC AUC; the area under a curve (called the ROC curve) that is created by plotting true positive rate vs. false positive rate at various thresholds. A ROC AUC of 0.5 is no better than chance and 1 is a perfect model, we achieved a ROC AUC of 0.680 for live birth prediction at blastocyst stage.

Next, we wished to compare our results against highly trained embryologist selection performance. To do this we used a subset of embryos (141 in total) for which Gardner grades^25^ (a set of ICM, TE and expansion scores standardly used by embryologists when assessing blastocysts) were available to produce ROC curves using the average model confidence scores and embryologist grades. In order to produce the ROC curve from embryologist grades we first converted the Gardner score letter grades into numbers, as proposed by Alpha Scientists in Reproductive Medicine and ESHRE Special Interest Group of Embryology^26^. Then an overall score was obtained by calculating an average of TE, ICM and expansion scores.

The ROC curves for both blastocyst assessment methods are shown in Fig. 2. The ROC AUC was similar for embryologist and model assessment, at 0.720 and 0.726 respectively. This suggests our model had a very similar performance to highly trained embryologist grading, as St Marys’ clinic host the UK NEQAS (a charitable consortium of external quality assessment laboratories) in reproductive biology. It is, however, important to note that the ML blastocyst model was at a disadvantage due to the use of limited information; only a single focal plane from just the final frame was used to train the ML model. In contrast, the assessment of the embryologist was done using multiple focal points from the final frame, with possible adjustment of the score according to the time-kinetic data.

**Figure 2:**
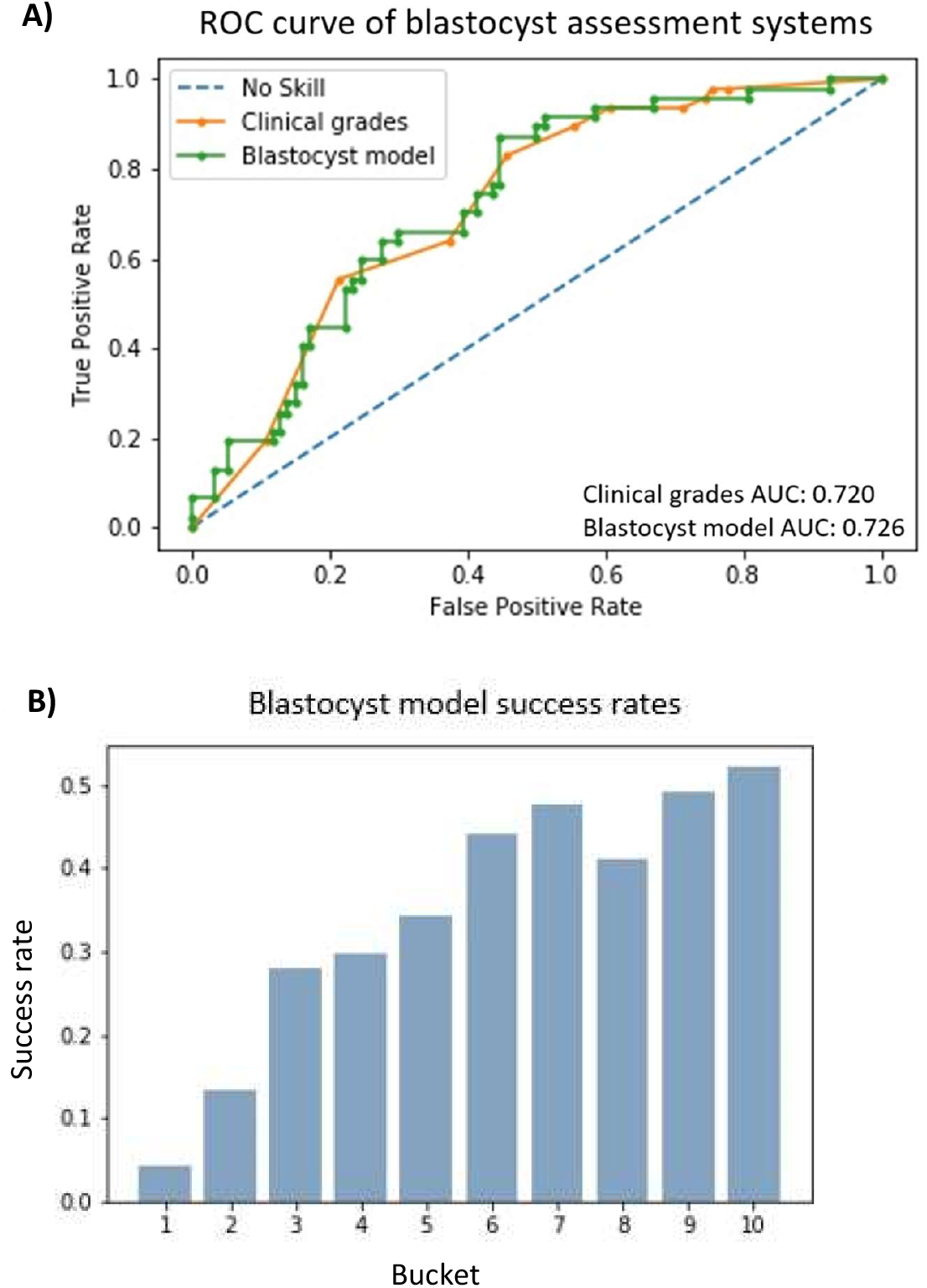
Live birth predicted from blastocyst stage. The blastocyst model predicts live birth using the last frame of the time-lapse video on day 5, the predictions produced by this model are a number between 0 and 1, 0 corresponding to an unsuccessful transfer (no pregnancy) and 1 corresponding to a successful transfer (resulting in live birth). The model scores used here are an average over 50 models (using 5-fold cross validation and 50 training attempts for each fold). **A)** A comparison between the performance of average model score and embryologist grading. Only embryos for which clinical grades were available were included, a total of 141. Clinical grades were originally based on the Gardner scoring system, these were then converted to numerical grades to allow a ROC curve to be generated. **B)** The correlation between live birth success rate and average model score, using cross-validation test scores for the whole dataset (700 embryos). The embryos have been sorted into buckets from 1 to 10 based on average model score, bucket 1 is the lowest scoring embryos and bucket 10 is the highest scoring embryos. The success rate of each bucket is the fraction of embryos in the bucket that resulted in a live birth.

In clinical practice, a numerical scoring system of embryo viability is more useful than a binary ‘successful’ or ‘unsuccessful’ prediction, as it allows the embryos in a cohort to be ranked in order of likelihood of live birth. Although our model was trained as a binary classifier, the confidence score is continuous. Confidence calibration varies between modelling systems^27^, therefore it was necessary to determine whether our model confidence scores could be useful in ranking embryos. To investigate this, we sorted all the embryos in our dataset into buckets by average test set model score, bucket 1 being the lowest score and bucket 10 being the highest score. The success rate (number of successful embryos in bucket/total number of embryos in bucket) for each bucket is shown in Fig. 2B. There was a general increase in success rate from bucket 1 to 10, suggesting that the confidence score was correlated to chance of live birth and therefore could be useful as a method to rank embryos in a cohort. However, increase in success rate with bucket number appeared to plateau from bucket 7-10, suggesting that there is a subset of high quality blastocysts (HQB) for which any further increase in model score has little correlation to increased chance of live birth. Analysing embryologist grade versus live birth for a dataset of fresh blastocyst transfers from St Marys we saw a similar plateau (Fig. S2A), suggesting that this is an issue for both ML and embryologist based approaches.

Our data demonstrated that it is possible to train MobileNetv2 to predict live birth from blastocyst morphology to a performance similar to embryologist assessment, with potential for further improvements to performance. Importantly, using both the ML and embryologist scores we have identified a limitation to embryo assessment based on blastocyst images; above a certain quality threshold it appears to be difficult to discriminate further the most viable embryo from a group.

### Identification of developmental time-points most indicative of viability

Currently, embryo assessment is based predominantly on the morphology of the embryo at the blastocyst stage. To establish whether earlier stages can be also be useful in assessing embryo viability, we extracted frames from each time-lapse video in the dataset at four pre-blastocyst stages that could be precisely defined; one hour before NEBD completion, first appearance of 2 cells, first appearance of 4 cells, and initiation of 8-16 cell division round, we referred to these stages as PN, 2-cell, 4-cell, and 8-16 cell respectively. Subsequently, live birth prediction models were trained using each stage.

As predicting live birth from embryo morphology alone is a difficult task, we decided to use our multi-class stage classification model as an extra domain-specific transfer learning step (see methods section for full details). We experimented with various hyper-parameters, the results for all stages are shown in Fig. S3A. Our results suggest that this domain-specific transfer learning step generally improved model performance, particularly for the PN and 4-cell models.

The ROC AUC values reported in Fig. S3A were the average of 50 repeat training runs each with a different randomly selected test set, as we had observed that variation due to the training uncertainty was minor compared to variation due to changes in the test set. This also allowed for a quicker investigation than the 5-fold cross validation with 50 training attempts used in the previous section. We chose to use varying test sets to increase statistical power as we found that separate test sets had a higher than desirable variability in performance, reflecting the heterogeneity of the data in this ML task (Fig. S3b). However, the transfer model was trained on the same dataset, so to check this was not resulting in an unfair bias it was necessary to also develop all models from scratch with a true ‘hold out test set’ – a test set separated out at the very start, prior to training the transfer model. As shown in table 1, our results continue to hold up in this control experiment, so going forward we use extra domain-specific transfer learning (with our optimised hyper-parameters) for the PN and 4-cell models in addition to the transfer learning from ImageNet (via the pre-trained layers) used for all other stages.

**Table I:** Performance on a hold out test using either the original MobileNetv2 model with fixed features up to the last layer or the MobileNetv2 model with fixed features and extra transfer learning; an extra hidden layer before the last layer has been pre-trained as a stage classifier. The number of hidden units in the last layer is the amount we found to be optimal for that developmental stage. ROC AUC scores shown here are the average scores on the test set over 50 training iterations. The test set contained 50 embryos in the successful class and 96 embryos in the unsuccessful class. The accompanying error bars are the standard error across these 50 training attempts.

| Stage | ROC AUC when standard MobileNetv2 model used | ROC AUC when extra transfer learning used |
| --- | --- | --- |
| PN | 0.547 +/- 0.0034 | 0.557 +/- 0.0023 |
| 2 cell | 0.555 +/- 0.0044 | 0.520 +/- 0.0042 |
| 4 cell | 0.533 +/- 0.0043 | 0.578 +/- 0.0043 |
| 8-16 cell | 0.578 +/- 0.0043 | N/A |
| Blastocyst | 0.714 +/- 0.0019 | 0.703 +/- 0.0016 |

The average test set ROC AUC over 50 training attempts with random train/validation/test split is shown in Fig. 3B for each stage. The blastocyst stage has the most obvious morphological differences between classes, with unsuccessful embryos less likely to form expanded blastocysts, and as expected this model had the best performance. However, the pre-blastocyst models still gave above chance predictions. From these results it was clear that different developmental stages carry different predictive power. To identify specific moments in development for which live birth predictions are the most successful we tested multiple time-points at regular intervals using each of the previous developmental stages as reference points (Fig. 3C-G).

**Figure 3:**
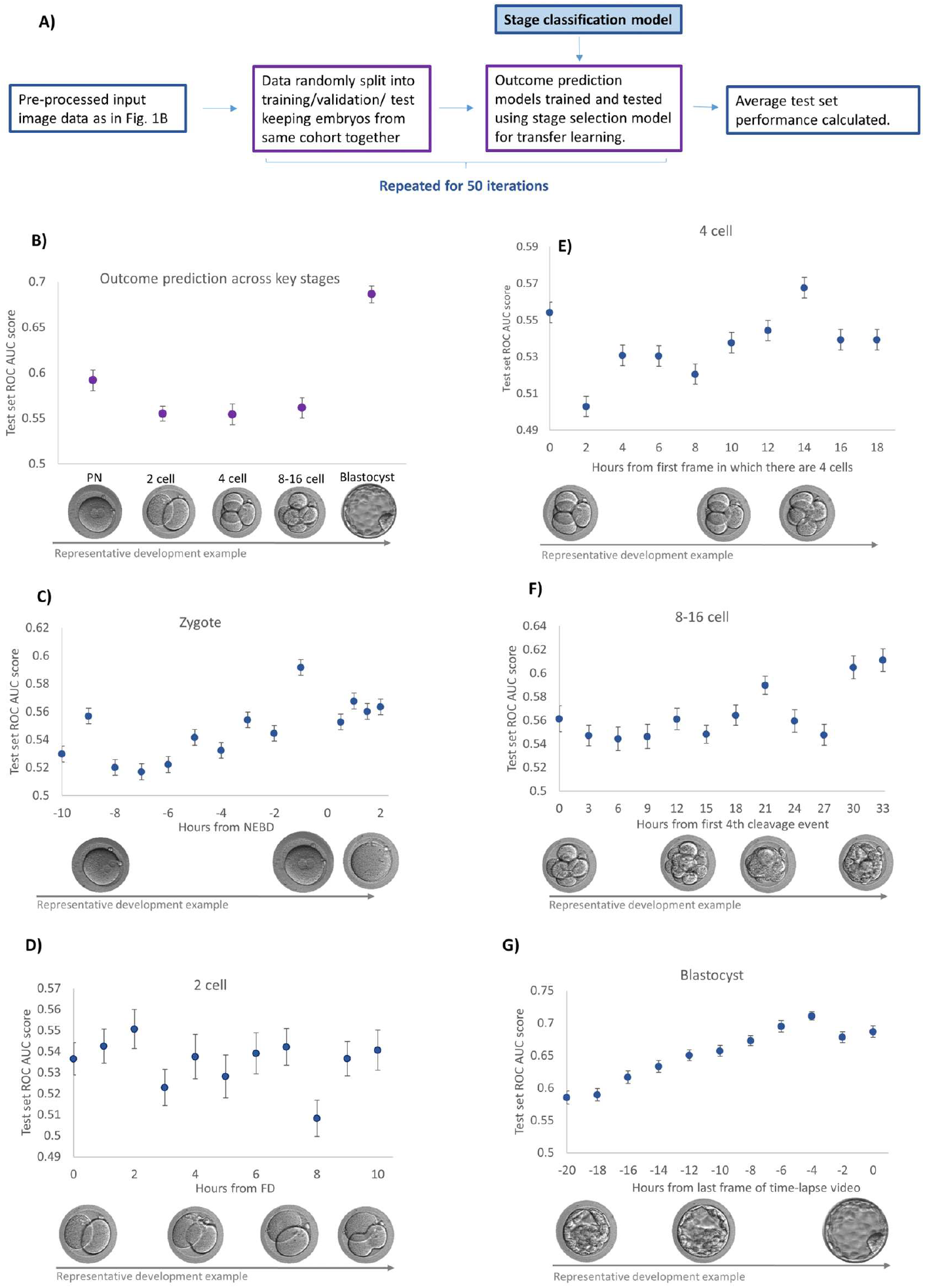
Predicting Live birth from a transferred embryo using specific time-points across development. All ROC AUC scores shown here are the average ROC AUC scores on the test set after 50 training iterations each with a different randomly allocated train/validation/test split with 25 embryos in the successful class and 48 embryos in the unsuccessful class for both the test and validation set. The accompanying error bars are the standard error across these 50 training attempts. A) The methodology for developing the live birth prediction models B) The ROC AUC scores for models trained to predict live birth using images from precisely defined moments of development. ‘PN ‘is one hour before completion of NEBD, ‘2 cell’ is first frame with 2 cells, ‘4 cell’ is first frame with 4 cells, ‘8-16’ cell is the first 4^th^ cleavage event, and ‘blastocyst’ is the last frame of the video. C) The ROC AUC scores for models trained to predict live birth using images at various time intervals before and after NEBD. D) The ROC AUC scores for models trained to predict live birth using images at various time intervals after FD. E) The ROC AUC scores for models trained to predict live birth using images at various time intervals after the first frame with 4 cells. F) the ROC AUC score of the test set for models trained to predict live birth using images at various time intervals after the first 4^th^ cleavage event. G) The ROC AUC scores for models trained to predict transfer live birth using images at various time intervals before the last frame of the time lapse video.

We found that the model performance varied at different time-points and appeared to peak at certain moments in development as shown in graphs 3C-G. For the PN stage (Fig. 3C), a performance peak was observed just before NEBD. Performance across the 2-cell stage (Fig. 3D) was variable with no peak seen. A performance peak was observed 14 hours after 4-cell (Fig. 3E), this generally is within the 4-8 cell transition however no potential explanation for this peak was found from visually examining the time-lapse videos. Another performance peak was observed 21 hours after initiation of 8-16 cell cycle (Fig. 3F), when embryos tend to be in the morula stage. As an initial examination of the time-lapse videos suggested that this was generally just before cavitation, we then quantified this by counting the number of cavitating embryos at this time-point and at time-points just before and after. The results, shown in table 2, confirmed that this performance peak corresponds to the moment just before cavitation. Lastly, the blastocyst stage showed a gradual improvement in model performance towards the end of the time-lapse video (Fig. 3G), appearing to reach a plateau 6 hours before the last frame, by which point the successful embryos have usually formed an expanded blastocyst. We found that the peaks at PN, 4-cell +14hrs and 8-16 cell +21hrs were all statistically significant when compared to a time-point 6 hours earlier (p values of 0.0001, 0.0005, and 0.0011 respectively).

**Table II:** Percent of embryos that have begun cavitating in each group at various time-points after first 4^th^ cleavage event.

| Hour | Percent of embryos cavitating |  |
| --- | --- | --- |
|  | Successful | Unsuccessful |
| 21 | 7.3% | 4.8% |
| 24 | 27.4% | 21.9% |
| 27 | 50.9% | 40.0% |
| 30 | 81.2% | 50.8% |

Finally, to test the potential benefit of averaging over repeat training attempts and also to obtain a single prediction for each embryo in our dataset (to be used in further investigations reported below), we re-trained each model using 5-fold cross validation with 50 repeat training runs to calculate an average test set model score for each embryo as in the previous section. The PN, 2-cell and blastocyst stage models were trained using the original time-point and the 4-cell and 8-16 cell stage models were trained using the peak performance time-point (4-cell +14hr and 8-16 cell +21hr). The ROC AUC scores calculated from these average model scores (table 3) showed that slightly better predictions can be made when the model score assigned to an embryo is averaged over many separately trained models, which is consistent with the existing literature on ensemble learning^28^. Additionally, we then also used the embryo average scores to calculate F1 and precision-recall AUC for each model (table S1) for further validation.

**Table III:** Effect of averaging score from many models rather than using just one model. The Average Individual ROC AUC scores are the average scores on the test set after 50 training iterations each with a different randomly allocated train/validation/test split with 25 embryos in the successful class and 48 embryos in the unsuccessful class for both the test and validation set. The accompanying error bars are the standard error across these 50 training attempts. For the ROC AUC using average confidence score, the average confidence score was first calculated from 50 models (using 5-fold cross validation and 50 training attempts) and these average confidence scores were then used to calculate the ROC AUC.

| Stage | Average Individual ROC AUC | ROC AUC using average confidence score |
| --- | --- | --- |
| PN | 0.591 +/- 0.0115 | 0.603 |
| 2 cell | 0.555 +/- 0.00779 | 0.575 |
| 4 cell | 0.567 +/- 0.0114 | 0.581 |
| 8-16 cell | 0.585 +/- 0.00779 | 0.585 |
| Blastocyst | 0.686 +/- 0.00937 | 0.680 |

### Combined model outputs allow refined ranking of high grade blastocysts

We next wanted to investigate whether information from the identified developmental time-points could provide an insight into the HQB group identified previously and assist in embryo selection. To test this, we calculated the ROC AUC of models at each stage on just the HQB group. The results (shown in Fig. 4A) show that the blastocyst model performed almost no better than chance and worse than all the earlier stage models, and the PN model performed the best. This suggests that if multiple embryos in a cohort fall into this HQB group (blastocyst model score>0.83), then pre-blastocyst stage models should be used to choose which one of these HQB embryos to transfer, rather than simply choosing the one with the highest blastocyst score.

**Figure 4:**
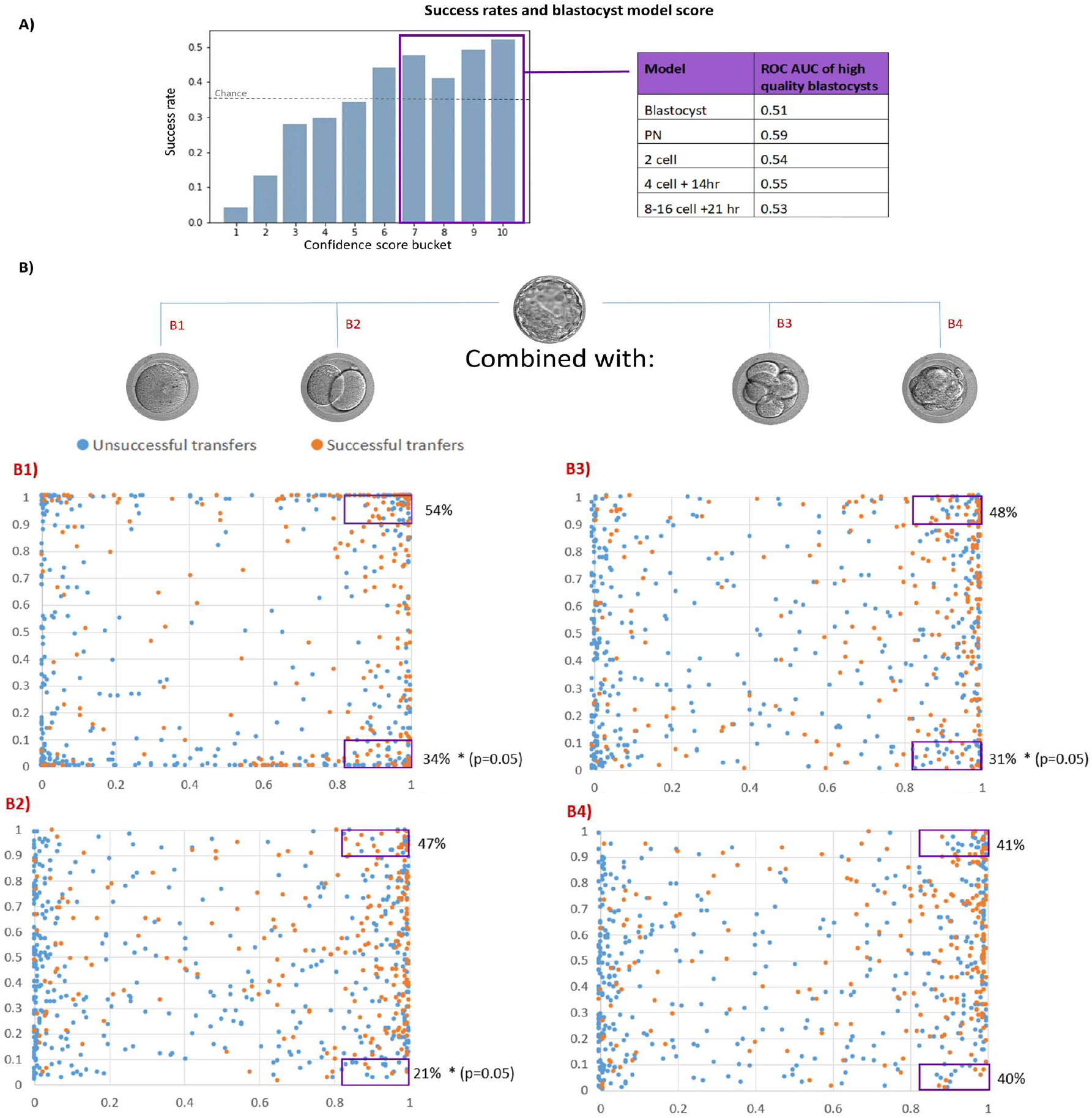
Ranking Embryos by blastocyst model score and combining model outputs. A) The association between live birth success rate and blastocyst model prediction (a model trained to predict live birth from images taken at blastocyst stage). The embryos have been sorted into buckets by blastocyst model confidence score, bucket 1 is lowest and bucket 10 is highest. The graph (left) shows the success rate of each bucket, defined as the fraction of embryos in each bucket that resulted in a live birth. The table on the right shows the ROC AUC for embryos in buckets 7-10 (“high quality blastocysts” – 259 in total) obtained using each of the models trained in this study. B1-4) Blastocyst model score vs. PN, 2 cell, 4 cell +14hr, and 8-16 cell+ 21hr model score respectively. The probability of live birth for high quality blastocysts (see A above) with a very high pre-blastocyst model score (>0.9) or very low pre-blastocyst model score (<0.1) is also shown for each graph.

To further investigate the benefit of using earlier stage models in conjunction with the blastocyst model we plotted the pre-blastocyst score vs. blastocyst score for each embryo, as shown in Fig. 4B. These graphs provide a visualisation of the spread of scores assigned by each model and the correlation between earlier stage scores and blastocyst score. We noticed that the model confidence scores had a tendency to be close to the extreme values; 0 or 1. To further investigate the added value of the pre-blastocyst stage models we calculated the success rate of the HQB embryos when the pre-blastocyst model score was >0.9 vs <0.1. This was then compared with the success rate of the HQB group as a whole (0.47) to calculate significance. We found that the success rate was significantly (p=0.05) lower when the PN, 2-cell, or 4-cell+14hrs model score is < 0.1. This suggests that the early models are most useful at identifying low quality embryos in the HQB group. This adds further evidence that there are developmentally abnormal embryos in the HQB group and pre-blastocyst model predictions should be taken into account when selecting from this group.

## Discussion

A better understanding of the processes driving human preimplantation development can help with embryo selection for IVF and assure that the best quality embryos are chosen for the transfer. Here, we identified previously unreported windows of development that are most closely linked to embryo viability and demonstrated that predictions from these developmental stages can be used to refine the selection for transfer of high quality blastocysts. Embryo assessment, both manual and ML, typically has an emphasis on specific chosen time-points, however these may not necessarily be the optimal moments of embryo development for assessing viability. We found the performance of live birth predictor models varied considerably depending on the exact moment in development in which they were trained, with peaks in performance found at multiple developmental stages. This could possibly be part of the reason for disagreements in the findings of studies linking embryo morphology to viability,(e.g the effect of vacuoles at zygote stage^29,30^) as it is possible researchers do not always use the exact same moments in development. Our findings could inform future embryo viability studies, as we have provided guidance on which developmental time-points to focus on. This could be particularly valuable for the morula, as there has been limited research into this stage yet our research suggests that during a specific window it may contain important information about embryo viability. Additionally, the plateau in performance found at the blastocyst stage could be of clinical relevance as it suggests that longer culture of the blastocyst beyond this point does not result in increased accuracy in assessment.

We noticed that the peaks in model performance found at PN stage and 8-16 cell +21hrs are both just before certain well described developmental events; NEBD and blastocyst cavitation respectively. There are two possible explanations for this; at these pre-event time-points there is less natural variation in the appearance of viable embryos (compared to time-points where the embryo is undergoing processes such as PN growth or compaction) so it is easier to distinguish important developmental abnormalities, or these may be biologically important moments where any deviation from normal development can prevent the embryo from developing properly. For example, the ability of embryos to correctly prepare and execute the first mitosis is one of the defining moments of development and any anomalies around this time, such as problems with the NEBD process, can result in the developmental failure. The peak during the transition from the 4 to 8 cell stage coincides with embryonic genome activation in human embryos^31,32^, raising the possibility that there are some morphological manifestations that can indicate the successful activation of the genome.

Finally, we investigated whether the previously identified key moments of development could help to understand the nature of the HQB group – a sub-group of embryos that all had a blastocyst model score above a certain threshold where any further increase in model score seemed to be unrelated to further increase in viability. It was not clear whether the embryos in this group were all of high quality and just failing for non-embryonic (e.g uterine) reasons or if there were embryos within the group that had developed abnormally despite receiving a high blastocyst model score. Our findings suggested the latter is true, as we found that it was possible to use the pre-blastocyst models to identify HQB embryos that were less likely to result in live birth. This provides evidence that predictions from early development may be used in conjunction with predictions from the blastocyst stage to give a better assessment of embryo quality than the blastocyst model alone. In addition to embryo selection it is also possible that the pre-blastocyst models could be used to help diagnose the reason for IVF transfer failure, if all HQB blastocysts in cohort fail to result in live birth then the earlier model predictions could provide an indication of whether there were embryonic issues not detectable at blastocyst stage or if the issue is more likely to be related to uterus.

Additionally, our findings could be used to support earlier transfers. Extending the culture until blastocyst stage is associated in some studies with a number of adverse outcomes, including pre-term birth, altered birthweight, mono-zygotic twinning, and shortened telomeres^33-37^. These disadvantages are not unexpected, since blastocyst culture acts to expose embryos to selection stress in a non-physiological in vitro environment, at the precise time in development when the embryonic genome and epigenome are being reset as part of the formation of a new individual^37-39^. Blastocyst culture is currently favoured as it leads to significantly higher live birth rate, per transfer and per embryo^40^, however the cumulative live birth rate, including transfer of all fresh and frozen embryos from a single egg collection event, are not necessarily increased using this policy. Our findings provide the flexibility of making viability assessments at various points in development, supporting various clinical strategies.

One of the biggest common limitations to ML is large data requirements. However, the models trained here demonstrate the potential of carrying out a CNN based analysis of a biological process without needing large amounts of training data. Our blastocyst model was competitive with highly trained embryologists and obtained a similar performance to other studies that used CNNs to predict live birth^20-22^, despite being trained on a single clinic dataset. This suggests that our algorithms could easily be re-trained on an individual clinics’ patient population to become specifically tailored to that clinic, an important factor as most clinics have distinct treatment policies and patient populations^41^ . The high performances achieved in stage classification also demonstrates the high capability of our chosen model on this limited dataset, even for quite subtle subcellular differences such as classifying embryos as before NEBD vs. after NEBD. Our stage classification models showed that increases in performance became small when training set size was increased beyond around 200-400, suggesting that the data amount was not likely to be the main limiting factor in performance.

The aim of this study was to provide a better understanding of pre-implantation development in the context of embryo assessment, therefore we focused on investigating the utility of various developmental stages rather than developing a new state of the art algorithm. It is quite possible that more advanced ML techniques could lead to improvements in performance, we hope the information provided here can inform future studies on embryo assessment via deep learning and contribute to improving embryo selection procedures before IVF.

## Conclusion

In summary, we have identified specific windows of early development that are most predictive of live birth by applying a readily accessible, pre-trained CNN (MobileNetV2) to hospital-specific image data. We have also provided evidence that information from the identified developmental time-points most indicative of viability could be combined with blastocyst stage model predictions to give better overall embryo assessment than selecting based solely on blastocyst morphology. Finally, the application to a single-clinic data set, of limited size, highlights the feasibility of exploiting state-of-the-art machine learning techniques in individual clinic settings, whilst being able to capture and cater for local protocols and data heterogeneity.

## Material and methods

### 1. Patients and time-lapse videos

Time-lapse videos of developing embryos were supplied by the IVF clinic in the Department of Reproductive Medicine at Old Saint Mary’s hospital, Manchester University NHS Foundation Trust, Manchester, UK. Embryos were cultured in either the Embryoscope ™ or Embryoscope+ ™ time-lapse incubator system (Vitrolife, Sweden). Throughout the study period there were no changes to the culture media or other culture conditions. Embryos were cultured in GTL overlaid with Ovoil (Vitrolife, Sweden) from post-injection (after ICSI, approximately 40 hours post-hCG trigger) to day 5 of development. Embryoscope ™ slides hold up to 12 embryos in individual wells holding 25µL of media, overlaid with 1.2ml oil. Embryoscope+ ™ slides hold up to 16 embryos, in 2 rows of 8 wells, each row of 8 wells is overlaid with 180uL of media. Embryos were not removed from the incubators during the observation periods, and media was not changed. Each time-lapse video exported had a framerate of 5-10 frames an hour. All images were irreversibly anonymised by clinic staff before being given to us.

The dataset comprised of fresh ICSI transfers from 2016-2019 that resulted in either live birth or no pregnancy, including both single embryo transfers (SET) and double embryo transfers (DET). Only DET that resulted in no pregnancy or male/female twins (to exclude the possibility of monozygotic twinning) were included. In total we used time-lapse videos for 443 successful embryos and 257 unsuccessful embryos.

### 2. Preparation of input image data

The frame number of specific stages in development were recorded by viewing each video in ImageJ. The frames at various time intervals before and after this moment were then automatically extracted using the timestamp (displayed in the bottom right of each frame). Measuring time in hours rather than frames was necessary as the time between frames was inconsistent between videos.

The datasets for each stage contained a frame from almost every embryo in the whole time-lapse dataset of 700 videos, however there were a small number of frames where the image quality was too low or the specific moment could not be determined (due to out of focus cell divisions or excessive fragmentation), therefore 10 embryos were excluded from the PN dataset, 13 from the 4 cell dataset, and 15 from the 8-16 cell dataset. We ensured the Embryoscope/Embryoscope plus ratio was equal across the two groups by randomly removing some of the successful Embryoscope embryos (this group initially had more embryos recorded by the EmbryoScope).

All images before the blastocyst stage were cropped to 300×300 pixels as this was the smallest size that captured the whole embryo including the zona. As embryos at the blastocyst stage expand a variable amount, occasionally almost filling the image, this stage was left uncropped. All images were then resized to 224×224 as this is the input size required by the pre-trained model.

### 3. Model

The model we have used is the MobilenetV2 model^42^, a CNN for which weights pre-trained on the ImageNet^43^ database are available. Using these weights allowed us to easily implement transfer learning from ImageNet, a standard strategy for relatively small datasets. MobileNetV2 is a computationally light model, therefore is convenient for any clinics wishing to repeat model training on their own dataset. We used fixed convolutional features, only training the final layer of the model. Prior to hyper-parameter tuning we used a base learning rate (BLR) of 0.0001, a drop out of 0.5 and the cross entropy loss function. For the live birth prediction models we applied a class weight of 2 to the live birth class to account for this class having about half the amount of data as the no pregnancy class.

For the live birth prediction models we also experimented with using developmental stage classification for extra domain-specific transfer learning. The first step was to add an extra hidden layer to the MobileNetV2 model and train it to predict developmental stage with both the hidden layer and last layer trainable (the rest of the network was fixed as before). For stages earlier than the blastocyst we used the classes as in Fig. 4E for step 1. For the blastocyst stage we trained a separate model with the before NEBD class replaced with a blastocyst class. In the 2nd step of training we then took this model as a starting point for our live birth prediction models, for this step the only trainable layer was the last layer, so all the convolutional layers and the hidden layer were fixed. We repeated these two steps with various numbers of hidden units (100,320,640, or 960) in the hidden layer to find the most suitable model structure.

### 4. Model training

Two training strategies were used in this work; cross validation, where train/validation /test sets were assigned randomly at the start of each repeat training run, and hold out test set, where training and validation sets were randomly assigned for each repeat run while the test set remained constant. We used a 80:10:10 train/validation/test split for the former and 60:20:20 train/validation/test split for the latter. When assigning embryos to the test and validation sets we kept embryos from the same cycle together for the live birth prediction models to avoid data leakage. We also kept the proportion of SET to DET constant throughout the train/validation/test sets. We then performed augmentation on the training set by rotating each image by 90,180 and 270 degrees and getting the mirror image.

Each training attempt was repeated twice, the first time we ran each model for 10,000 epochs and recorded at what epoch the validation set peaked in performance on average across all repeat training runs. The second time we trained the model for this optimal number of epochs for every repeat training run. We did this because our models were prone to overfitting so finding the optimal number of epochs was important, however the small size of our validation set led to a lot of fluctuation so averaging over many repeat runs was required.

### 5. Model performance evaluation

For each model setup we ran 50 repeat training runs and for each training run the area under the curve (AUC) of the receiver operator characteristic (ROC) was calculated. The ROC is a graphical plot of true positive rate vs. false positive rate at various thresholds. The AUC of this curve is therefore a measure of how well the model performs, 1 corresponding to a perfect classifier. We calculated the ROC AUC using Scikit-learn python library. The standard error in ROC AUC values over all repeat runs was then used to calculate error bar values.

The blastocyst model was compared to the embryologist scoring system using grades assigned by the embryologists at St Marys. An overall ‘embryologist score’ was calculated from averaging TE, ICM and expansion scores, we acknowledge this scoring system is imperfect as in reality the embryologist selecting the embryo may be able to select the better embryo out of embryos given the same grade and may place a different level of importance on the individual expansion/TE/ICM scores. However, due to the lack of an evidence base for selecting amongst embryos of similar grades we decided a simple average would be best for calculating an overall score, allowing for a comparison with our blastocyst model.

The comparison of model performance at the optimal time-point vs. the time-point 6 hours before (Fig.3) was carried out by a students’ t test. The significance between the success rates of subsets of the HQB group and the whole HQB group (Fig.4B) was calculated using a binomial test.

## Ethics

As this research used only fully anonymized, routinely collected data, ethical approval was not required in accordance with the NHS Health Research Authority (HRA) guidelines (http://www.hra-decisiontools.org.uk/ethics/”) Therefore written consent from participants was waived.

## Author contribution

B.P, D.B and J.H contributed to the concept of the study, H.H prepared the data, C.M pre-processed the data, developed the models and analysed the results. B.P, J.H and C.M designed the modelling pipeline. C.M drafted the paper and B.P, J.H, D.B and H.H critically reviewed paper.

## Data availability

The data that support the findings of this study are available from the corresponding author upon reasonable request.

The code used to build the models is available at https://github.com/CMapstone/eMLife.

## Acknowledgements

We would like to thank other members (current and past) of the Plusa lab for their discussion about the paper. C.M. was sponsored by the Wellcome Trust as part of the Quantitative and Biophysical Biology programme (220001/Z/19/Z). Funding to B.P Lab was provided by the Wellcome Trust grant Seed Awards in Science (212372/Z/18/Z). D.B was funded by Manchester University NHS Foundation Trust.

