## Supplementary Material for "Deep learning pipeline reveals key moments in human embryonic development predictive of live birth in IVF"

**Supplementary Data**

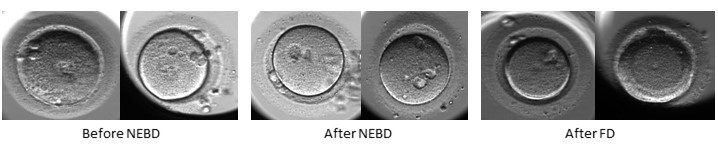

**Figure S1A: Examples of images incorrectly classified by stage selection models.** Left: images before NEBD where the PN are hard to see. Middle: images without PN with vacuoles that resemble PN. Right: 2 cell embryos that look like 1 cell embryos.

| **Bucket** | **Embryologist grades included** | **Number in bucket** |
| --- | --- | --- |
| 1 | Cavitating/morula | 5 |
| 2 | 1BB,2AB,2BB,2CC | 16 |
| 3 | 3BB, 3BC,3CB | 40 |
| 4 | 3AA,3AB,3BA | 37 |
| 5 | 4BB | 89 |
| 6 | 4AB 4BA | 157 |
| 7 | 4AA | 147 |
| 8 | 5BB | 36 |
| 9 | 5AB 5BA | 134 |
| 10 | 5AA | 182 |

Success rate vs Embryologist grade

| Bucket | Success rate |
| --- | --- |
| 1 | 0.2 |
| 2 | 0.142857 |
| 3 | 0.425 |
| 4 | 0.324324 |
| 5 | 0.370787 |
| 6 | 0.44586 |
| 7 | 0.428571 |
| 8 | 0.444444 |
| 9 | 0.470149 |
| 10 | 0.527473 |

**Figure S2a: Embryologist grade vs success rate.** From a datset of 843 fresh blastocyst transfers.

4 cell stage

Validation set ROC AUC

| Number of units in final layer | BLR 0.01 | BLR 0.001 | BLR 0.0001 |
| --- | --- | --- | --- |
| 100 | 0.529908 | 0.521019 | 0.51468 |
| 320 | 0.525673 | 0.542487 | 0.524193 |
| 640 | 0.523599 | 0.556256 | 0.568943 |
| 960 | 0.537169 | 0.536583 | 0.519948 |
| 1280 | 0.505757 | 0.527883 | 0.519038 |

8-16 cell stage

Validation set ROC AUC

| Number of units in final layer | BLR 0.01 | BLR 0.001 | BLR 0.0001 |
| --- | --- | --- | --- |
| 100 | 0.536586 | 0.504178 | 0.499582 |
| 320 | 0.501859 | 0.514339 | 0.510142 |
| 640 | 0.53349 | 0.545552 | 0.556402 |
| 960 | 0.560824 | 0.543699 | 0.548406 |
| 1280 | 0.542866 | 0.561377 | 0.592044 |

2 cell stage

Validation set ROC AUC

| Number of units in final layer | BLR0.01 | BLR 0.001 | BLR 0.0001 |
| --- | --- | --- | --- |
| 100 | 0.518654 | 0.547474 | 0.559832 |
| 320 | 0.53517 | 0.546449 | 0.547864 |
| 640 | 0.554087 | 0.52673 | 0.528165 |
| 960 | 0.522195 | 0.542407 | 0.545089 |
| 1280 | 0.534142 | 0.546823 | 0.547389 |

PN stage

Validation set ROCAUC

| Number of units in final layer | BLR0.01 | BLR 0.001 | BLR 0.0001 |
| --- | --- | --- | --- |
| 100 | 0.525879 | 0.549054 | 0.536079 |
| 320 | 0.541546 | 0.552825 | 0.544137 |
| 640 | 0.563421 | 0.591715 | 0.589382 |
| 960 | 0.527749 | 0.567964 | 0.560106 |
| 1280 | 0.525757 | 0.570142 | 0.560839 |

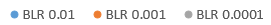

**Figure S3A**: **Hyper-parameter tuning for each stage; examining the effects of varying BLR and using extra domin-sprecific transfer learning.** Models with 100-960 units use an extra hidden layer that had been pre-trained on the stage classification dataset. The models with 1280 units use the standard MobileNetv2 model with no extra hidden layers.

Blastocyst

Validation set ROC AUC

| Number of units in final layer | BLR 0.01 | BLR 0.001 | BLR 0.0001 |
| --- | --- | --- | --- |
| 100 | 0.672341 | 0.652831 | 0.682019 |
| 320 | 0.670198 | 0.677785 | 0.673174 |
| 640 | 0.659812 | 0.675707 | 0.695286 |
| 960 | 0.664838 | 0.676731 | 0.675688 |
| 1280 | 0.657983 | 0.678181 | 0.68085 |

Blastocyst

4 cell

2 cell

PN

Comparing different test sets using original model (no extra TL)

ROC AUC

|  | Test set 1 | test set 2 | test set 3 |
| --- | --- | --- | --- |
| 1 | 0.541928 | 0.551361 | 0.584667 |
| 2 | 0.545576 | 0.514059 | 0.512444 |
| 3 | 0.543484 | 0.54898 | 0.518929 |
| 4 | 0.567598 | 0.574817 | 0.491366 |
| 5 | 0.700181 | 0.701877 | 0.712723 |

**Figure S3B:** **Comparison of the ROC AUC obtained by 3 different hold out test sets at each stage.** A BLR of 0.0001 and the original MobileNetv2 model with no extra domain-specific transfer learning was used.

|  | **F1, Live birth positive** | **F1, no pregnancy positive** | **Precision-recall AUC, live birth positive** | **Precision-recall AUC, no pregnancy positive** |
| --- | --- | --- | --- | --- |
| **PN** | 0.478 | 0.661 | 0.412 | 0.737 |
| **2 cell** | 0.452 | 0.558 | 0.385 | 0.560 |
| **4 cell** | 0.475 | 0.555 | 0.411 | 0.554 |
| **8-16 cell** | 0.468 | 0.561 | 0.405 | 0.570 |
| **Blastocyst** | 0.4639 | 0.6468 | 0.571 | 0.816 |

**Table S1: F1 and precision-recall AUC values for each model.** Each metric has been calculated twice, once with the live birth class labelled as positive and once with the no pregnancy class labelled as positive.
